# Characterization of a novel botybirnavirus with a unique dsRNA infecting *Didymella theifolia* from tea plants

**DOI:** 10.1101/2022.12.06.519410

**Authors:** Liangchao Ye, Xinyu Shi, Yunqiang He, Jiao Chen, Qingeng Xu, Karim Shafik, Wenxing Xu

**Author notes:** Corresponding author: Wenxing Xu, Professor, Plant Pathology.

## Abstract

*Didymella theifolia* specifically infects some local varieties of *Camellia sinensis* in China, representing a unique fungal species, and characterization of the mycoviruses related to this fungal species is attractive. Three double-stranded RNAs (dsRNAs; dsRNAs 1, 2 and 3 with sizes of 6338, 5910 and 727 bp, respectively) were identified in an avirulent strain CJP4-1 of *D. theifolia* exhibiting normal growth and morphologies. Characterization of the dsRNAs 1 and 2 revealed that they are genomic components of a novel botybirnavirus, tentatively named Didymella theifolia botybirnavirus 1 (DtBRV1), and encapsidated in isometric virions with a size of ∼39.8 nm in diameter. It is worth noting that dsRNA3 shares no detectable identity with those sequences deposited in NCBI database, while a high identity (36.58% and 40.93%) with the left regions of dsRNAs 1 and 2, but is not encapsidated in DtBRV1 particles, suggesting it is a unique dsRNA unit that is not a DtBRV1 component or a satellite and its taxonomic classification remains unclear. SDS-polyacrylamide gel electrophoresis in combination with peptide mass fingerprint analysis revealed that DtBRV1 capsid protein consisting of polypeptides encoded by the left regions of both genomic components. DtBRV1 is efficiently vertically transmitted through conidia while difficult in horizontal transmission from strain CJP4-1 to other strains. DtBRV1 has no effects on fungal growth and virulence as accessed with the transfectants of virulent strain JYC1-6 of *D. theifolia* infected by DtBRV1. DtBRV1 with specific molecular traits contributes useful information for a better understanding of the mycoviral community.

**Importance:** Tea plants represent an ancient and unique plant species community cultured in China, while the mycoviruses related to the phytopathogenetic fungi infecting tea remain limited. Here, we characterized a novel botybirnavirus (tentatively named Didymella theifolia botybirnavirus1 (DtBRV1), and a specific dsRNA infecting *Didymella theifolia* responsible for a noticeable disease of tea plants. DtBRV1 contains two dsRNAs (1 and 2) encapsidated in isometric virions in size of ∼39.8 nm, while dsRNA3 is not encapsidated in the viral particles although it has a high identity with the mycoviral genomic components. Additionally, DtBRV1 coat proteins are composed of fused proteins encoded by both dsRNA-coding open reading frames most likely after cleave and fuse processing progress, which is striking unlike most mycoviruses. With some specific molecular traits, DtBRV1 and the related specific dsRNA unit expand our understanding of virus diversity, taxonomy, and evolution.

## INTRODUCTION

Mycovirus (either termed fungal virus) has been reported widely in all major taxa of fungi including filamentous fungi, yeasts, and oomycetes (1-5) since it was first discovered from *Agaricus bisporus* (a kind of mushroom) in 1962(6). Based on the nature and function of nucleic acids, mycoviruses have been taxonomically grouped into 21 families and some unclassified taxa (7, 8) by the International Committee on Taxonomy of Virus (ICTV, https://talk.ictvonline.org/taxonomy/), including nine with positive-sense single-stranded RNA (+ssRNA) (*Alphaflexiviridae, Barnaviridae, Botourmiaviridae, Deltafleiviridae, Endornaviridae Gammaflexiviridae, Hypoviridae, Mitoviridae, and Narnaviridae*) (9), six (*Chrysoviridae, Partitiviridae, Quadriviridae, Reoviridae, Totiviridae, and Megabirnaviridae*) with double-stranded (dsRNA) genomes(10), one with negative-sense ssRNA (-ssRNA; *Mymonavirida*e) (7), one with ssDNA (*Genomoviridae*) (11). Many of these mycoviruses parasite their host fungi in a latent manner, while some can dramatically attenuate the virulence of host fungi, even change their lifestyle from a pathogen into an endophyte, and are candidates as biocontrol agents against fungal diseases (11-14), exemplified by successful control of chestnut blight by Cryphonectria hypovirus 1 (CHV1) in Europe (6).

Mycoviral numbers have largely increased in recent years with the aid of the next-generation sequencing (NGS) approach after this method was developed in the second half of the 2000s, whereas those associated with fungi infecting tea plants (*Camellia sinensis* (L.) O. Kuntze) are still very limited. Tea is an important beverage crop in tropical and subtropical regions, ranking as the most consumed beverage in the world after water in the past decades, and has gained favor with consumers as a healthy beverage (15). More importantly, tea plants originated from China and has been cultivated for over 3000 years, representing an ancient and unique plant species community in Asia, and characterization of the mycoviruses associated with tea plants would open a window to realize such a unique microbiological community. *Didymella theifolia* W.X. Xu and Y.Q. He is a newly identified and named fungal species infecting tea trees causing tea leaf brown-black spot disease, locally known in Chinese as “chixingbing”, characterized by tiny blackish- or reddish-brown spots on the tender leaves, and affects tea quality and causes it to taste bitter and astringent (16). No mycoviruses are identified and characterized from this fungal genus.

*Botybirnavirus* is a newly identified genus with the first member identified in 2012, while has not been classified into any families by the ICTV although the family Botybirnavirade has been proposed once upon a time when the first botybirnavirus was identified (17). Up to now, only six members belonging to this genus are identified and characterized in three phytopathogenic fungi, including *Botrytis porri, Sclerotinia sclerotiorum*, and *Alternaria* spp., and all have a bipartite genome consisting by two dsRNA components with similar sizes encapsidated in isometric virions(17-21). An RNA-dependent RNA polymerase (RdRp) domain is easily detected in an open reading frame (ORF) encoded by one dsRNA strand, while their coat proteins (CPs) remain undetermined. Overall, this mycoviral community remains far less characterized.

Here, a bipartite dsRNA mycovirus was detected in *D*.*theifolia*, and identified as a novel botybirnavirus based on the sequence alignment and phylogenetic analysis. This botybirnavirus presents some specific molecular and biological traits and will contribute useful information for a better understanding of this viral community.

## RESULTS

### Three dsRNAs in strain CJP4-1 of *D. theifolia*

Compared with *D. theifolia* strains JYC1-6 and JYC1-9, strain CJP4-1 showed similar colony morphologies except for the production of more dark pigments but no virulence on detached tea leaves, since no lesions were observed on the inoculated tea leaves for strain CJP4-1, while long lesions ranging from 8.0 mm to 17.0 mm for the reference strains (Fig. 1A, B, and C). To investigate whether a mycovirus was responsible for the attenuated virulence of strain CJP4-1, the mycelia of these strains were subjected to dsRNA extraction and agarose gel electrophoresis. While three dsRNAs (termed dsRNAs 1, 2, and 3 according to their decreasing sizes), resistant to the digestion with DNase I and S1 nuclease, were detected in preparations of strain CJP4-1, no corresponding dsRNA bands were observed in preparations from the other strains (Fig. 1D). The sequences of the full-length cDNAs of dsRNAs 1 and 2, in size of 6338 and 5910 bp respectively, were determined by NGS approach in combination with informatics analysis and RACE protocols, and dsRNA3 in size of 727 bp was obtained by amplification with PC2 primer after ligated with PC3-T7loop oligo and reverse transcribed (Fig. 2A).

**Fig. 1.**
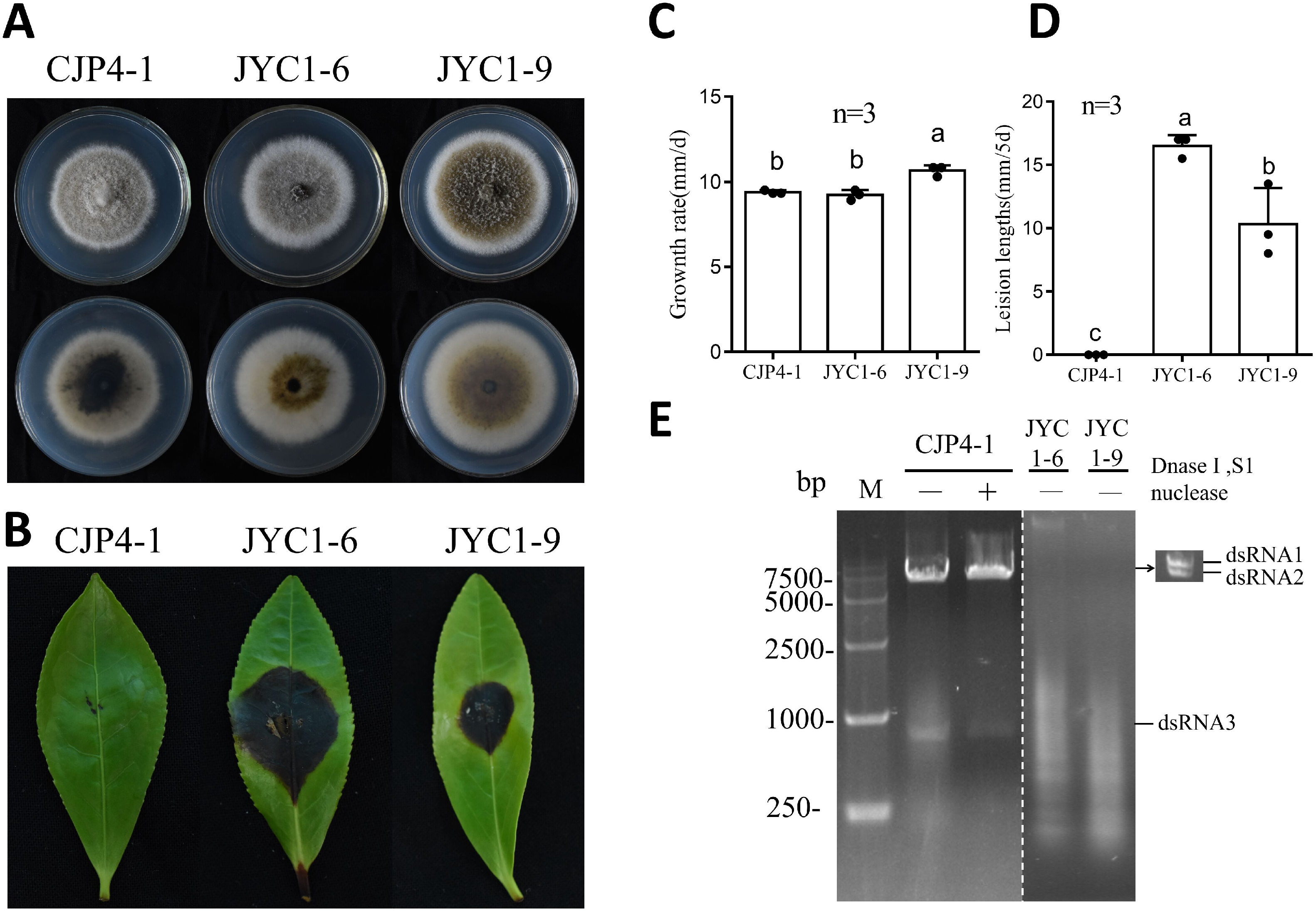
Morphologies, growth rates, virulence, and nucleic acid extraction of strains CJP4-1, JYC1-6, and JYC1-9 of *Didymella theifolia*. (A) Colony morphologies of *D. theifolia* strains cultured on PDA at 25°C for 7 days. (B) Virulence test of the *D. theifolia* strains on the detached tea leaves (*Camellia sinensis* var. E’cha no.1) under wounded conditions at 5 days post inoculation (dpi). (C and D) Bar graph for the growth rates (C) and lesion lengths (D) of *D. theifolia* strains. (E) Agarose gel electrophoresis of nucleic acids extracted from *D. theifolia* strains, without (-) or with (+) enzyme treatment by nuclease S1 and DNase I.

**Fig. 2.**
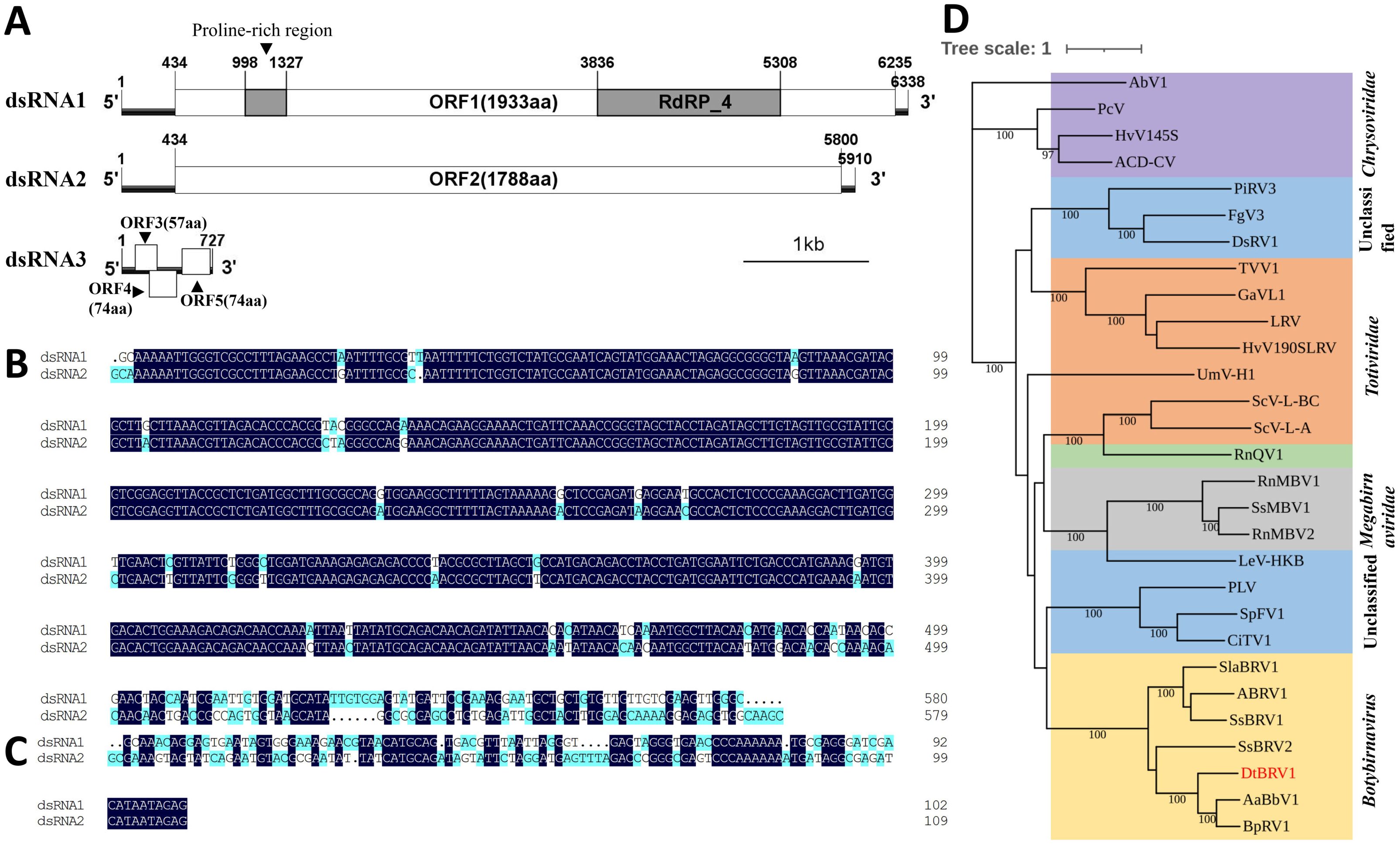
Genomic organization and phylogenetic analysis of DtBRV1. (A) A schematic diagram of the genomic organization of dsRNAs 1 to 3. The start and end positions are labeled for the genome, untranslated regions (UTR), the open reading frame (ORF), and an RNA-dependant RNA polymerase (RdRp)-4 superfamily domain. (B and C) Alignment of the 5-terminal and 3-terminal sequences of the coding strands of both dsRNAs-1 and 2, respectively. Identical nucleotides are highlighted with dark blue color. (D) A maximum likelihood phylogenetic tree was constructed using IQ-TREE software based on RdRp sequences of DtBRV1 and the related dsRNA viruses. The bootstrap values were deduced based on 1000 replicates, and bootstrap values lower than 70% are not shown in the figure. The scale bar refers to the genetic distance of 1. Definitions, abbreviations, and other related information of the referred viruses are listed in Table 1.

### DsRNAs 1 and 2 compose the genome of a novel botybirnavirus

The 5’-untranslated regions (5’-UTRs) of both dsRNAs 1 and 2 are in the same size of 433 bp, and share a high sequence identity of 94.24% (Fig. 2B). The sizes of the 3’- UTRs of dsRNAs 1 and 2 are 103 and 110 bp, respectively, and the sequence identity is 54.05% (Fig. 2C).

**Table 1.**
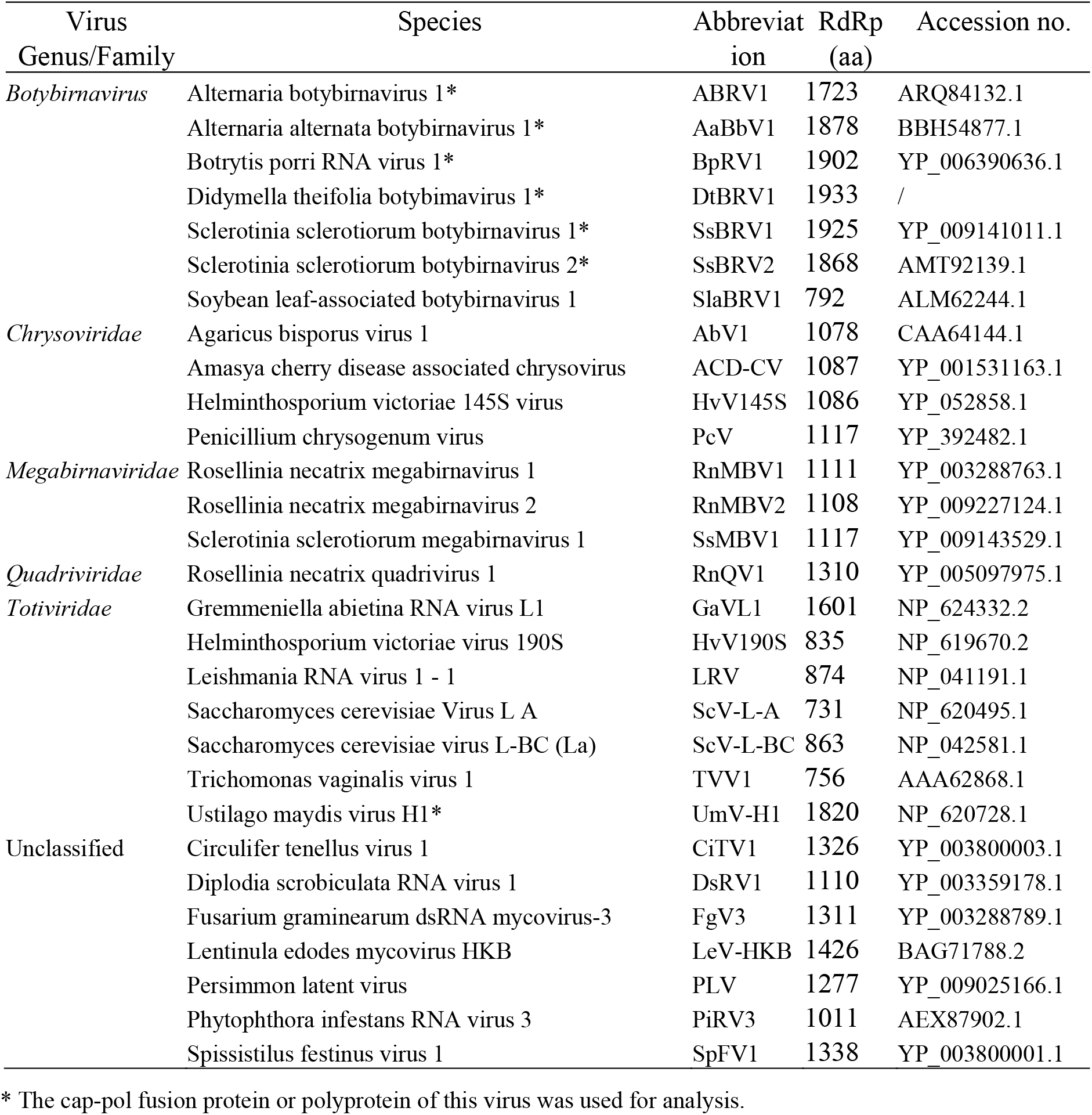
The infomation for the viruses used for phylogenetic analysis

Both dsRNA1 and dsRNA2 were predicted to encode an ORF, termed ORF1 (at nt positions 434-6235) and ORF2 (434-5800), respectively. ORF1 encodes a polypeptide in size of 1933 aa with an estimated molecular mass of 217.3 kDa, and contains a proline-rich domain at bp positions 998 to 1326 and an RNA-dependent RNA polymerase (RdRp)_4 superfamily structural domain at 3836-5308, with an estimated molecular mass of 56.5 kDa. The predicted RdRp sequence harbors eight conserved motifs (I-VIII), similar to those of other dsRNA viruses (Fig. S1). BLASTp searches unveiled that the ORF1-encoding polypeptide had the highest identity to the polypeptide encoded by Sclerotinia sclerotiorum botybirnavirus 3 (SsBRV3, accession no. AWY10943.1, *E-value* 0.0, coverage 98%, identity 46.98%). The maximum likelihood phylogenetic tree based on the RdRps from the related dsRNA viruses (Table 1) revealed that it was clustered together with the members of genus *Botybirnavirus*, and most closely phylogenetically related to Alternaria alternate botybirnavirus 1 (AaBRV1) and Botrytis porri botybirnavirus 1 (BpRV1) (Fig. 2D). ORF2 encodes a putative protein in size of 1788 aa with an estimated molecular mass of 198 kDa, and has high identities to those SsBRV3, BpBRV1 and AaBRV1 (*E-value* 0.0, coverage 95-99%, and identity 39.86-40.37%) whose function remains unknown. These results suggest that dsRNAs 1 and 2 are the genomic components of a novel botybimavirus, which is tentatively named Didymella theifolia botybimavirus 1 (DtBRV1).

For dsRNA3, three small ORFs (ORF 3 to 5) were predicted (with two in the plus strand and one in the minus) to encode three putative peptides in size of 57, 74, and 74 aa with estimated molecular masses of 6.4, 8.2 and 8.3 kDa, respectively, which are most likely not *bona fide* proteins due to so mall sizes. BLASTp searches unveiled that these putative peptides had no significant identity to known proteins. Similarly, BLASTn and BLASTx searches unveiled that dsRNA3 had no significant identity with those sequences deposited in NCBI. However, the sequence alignment of dsRNA3 with dsRNAs 1 or 2 showed that it has an identity of 40.93% with dsRNA1 at nucleotide positions 57-796 bp, and 36.58% with dsRNA2 at 57-806 bp (Fig, S2). The dsRNA3 was lost during the purification of virus particles and vertical transmissions (Fig. 3A and 4A), and even sometimes in the parent strain CJP4-1 after subcultures. Therefore, we speculate that the presence of dsRNA3 is unstable and not a component of virus DtBRV1.

**Fig. 3.**
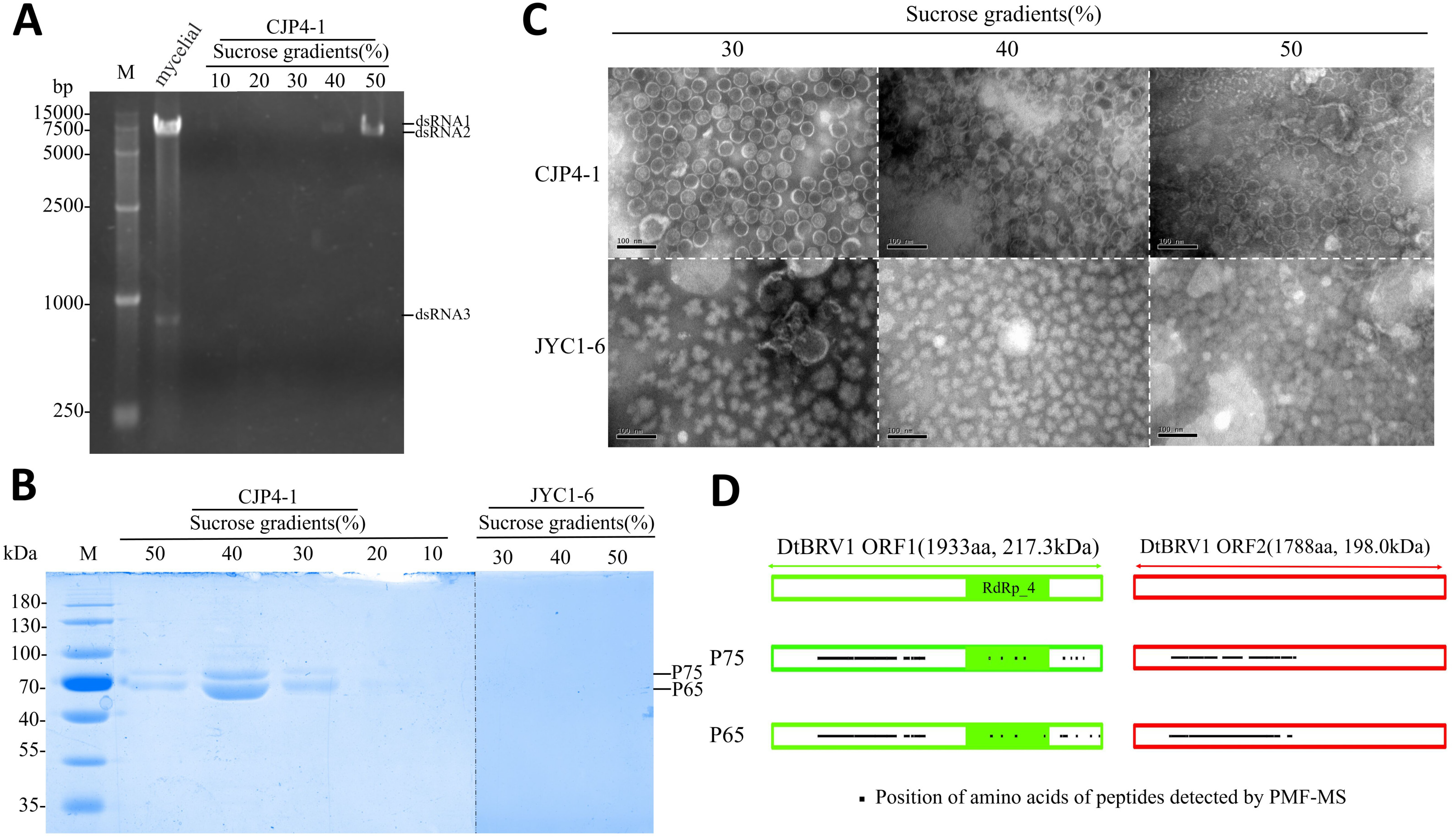
Virus-like particles, dsRNAs, and proteins extracted from the sucrose fractions following sucrose gradient centrifugation. (A) Agarose gel electrophoresis analysis of dsRNAs extracted from mycelia of strain CJP4-1 and purified virus-like particles of DtBRV1 from 10 to 50% sucrose fractions at 10% increments. M, DNA size marker. (B) SDS-PAGE analysis of the proteins from strains CJP4-1 and JYC1-6 in 10–50% sucrose fractions (with 10% increments) after sucrose gradient centrifugation. M, protein molecular weight marker. (C) TEM observation of VLPs of strain CJP4-1 and preparations from DtBRV1-free strain JYC-1-6 in 30 to 50% sucrose fractions. (D) Peptide mass fingerprinting analysis of proteins in purified virus particle fractions. Predicted coding regions for p75 and p65 are shown in the diagram for the DtBRV1 ORFs. The green box in ORF1 shows the position of the RdRp_4 domain. Dots in boxes indicate the positions of peptide fragments detected by PMF-MS (Tables S2 and S3).

**Fig. 4.**
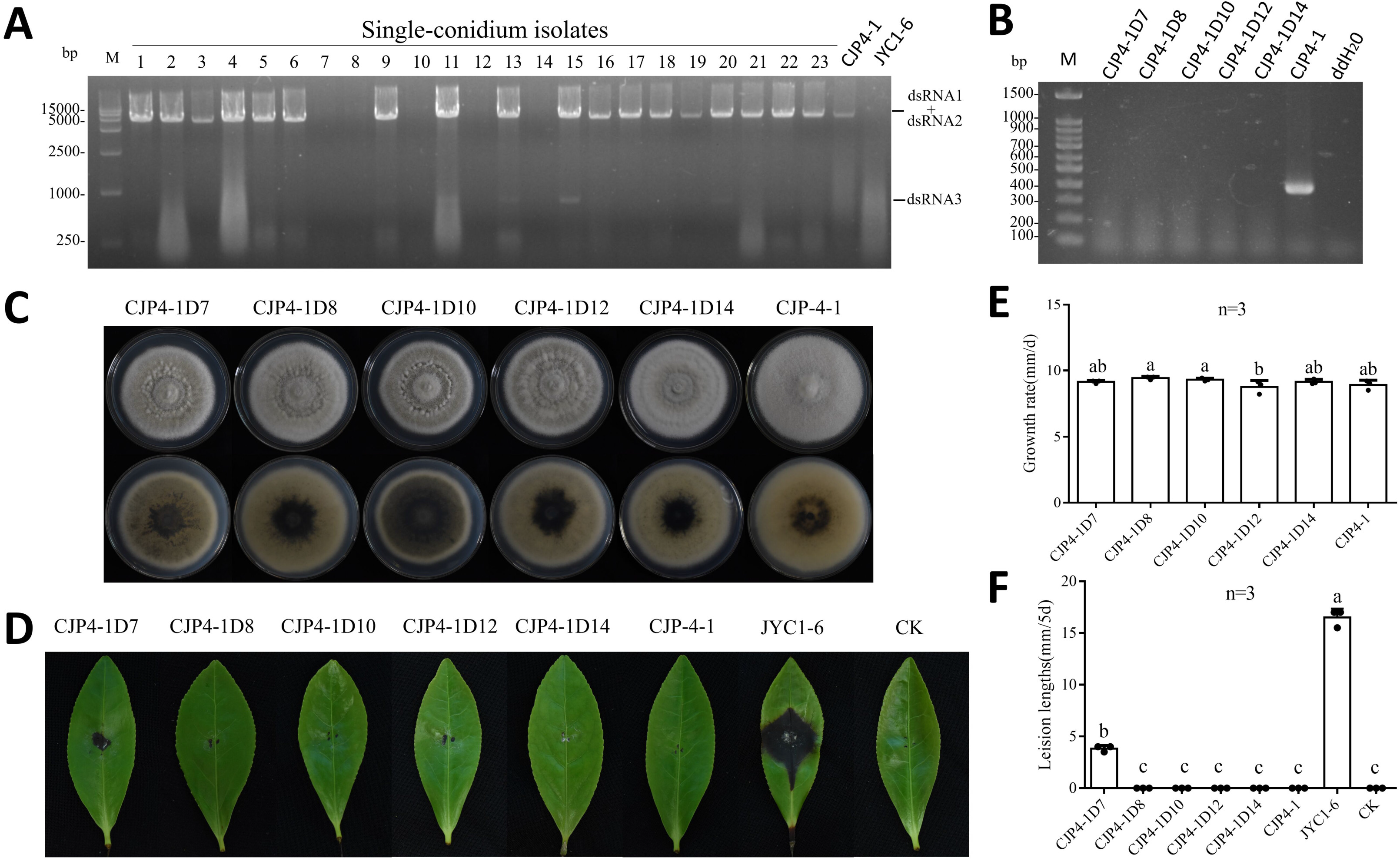
Transmission of DtBRV1 and effects on its host. (A) DsRNA extraction from the mycelia of single-conidium generated subisolates derived from strain CJP4-1. Lanes 1 to 23 are single-conidium isolates (CJP4-1D1 to CJP4-1D23). Strains CJP4-1 and JYC1-6 were involved as positive and negative controls. (B) RT-PCR identification of the DtBRV1 absence in subisolates CJP4-1D7, -D8, -D10, -D12, and -D14, with strain CJP4-1 involved as a positive control resulting in the target product in size of 390 bp, and double distilled water (ddH_2_O) as blank control. (C) Colony morphologies of strain CJP4-1 and its subisolates that DtBRV1 was eliminated cultured on PDA at 25°C for 7 days. (D) Virulence assessment of strain CJP4-1 and its subisolates on tea leaves (*C. sinensis* var. E’cha no.1). Virulent strain JYC1-6 was used as positive controls, and uncolonized PDA plug as black control (CK). (D and E) Bar graphs of the growth rates (D) and lesion lengths (E) of strain CJP4-1 and its subisolates.

### DsRNAs 1 and 2 are encapsidated in isometric virions

Viral particles (VLPs) of strain CJP4-1 were purified by centrifugation in stepwise sucrose gradients (10-50%, sucrose increments of 10%). Nucleic acids and proteins were extracted from each sucrose gradient fraction by ultracentrifugation, and subjected to electrophoresis analysis on agarose gel and SDS-PAGE, respectively. The results showed that both nucleic acids and proteins were mainly distributed in the 30-50% sucrose fractions (Fig. 3A and B). By the SDS-PAGE, four protein bands were observed on the gel between the molecular weight sizes of 100 and 60 KDa, with both bigger ones remaining faint while both smaller ones highly concentrated, while these proteins were not detected in the preparations from strain JYC1-6 (Fig. 3B). Examination of the gradient fractions by transmission electron microscopy (TEM) revealed isometric virus-like particles with a diameter of ∼39.8 nm in the 30-50% sucrose fractions, while these isometric virus-like particles were not observed in the preparations from DtBRV1-free strain JYC1-6 (Fig. 3C).

### DtBRV1 structure proteins are joint peptides encoded by both dsRNAs 1 and 2

To further verify the presence of the deduced proteins on SDS-PAGE analysis, those two dominant protein bands, tentatively termed p75 and p65 as referring to their approximate sizes, were separately excised from the gel and subjected to peptide mass fingerprinting (PMF) analysis. A total of 197 peptide fragments from p75 matched the partial sequence of ORF1-encoding polypeptide, accounting for 32% of the entire polypeptide, dominating at aa positions 267-891 (with 189 peptide fragments), while 271 matched the partial sequence (at aa positions 214-931) of ORF2-encoding protein, accounting for 34% of the entire coverage; 318 peptide fragments from p65 matched the partial sequence of ORF1-encoding peptide, accounting for 34% of the entire coverage, dominating at aa positions 267-897 (with 189 peptide fragments), while 271 matched the ORF2-encoding protein at aa positions 202-907, accounting for 36% of the entire coverage (Fig. 3D, and Tables S2 and S3). These results suggest that both p75 and p65 are dominantly derived from both left regions of ORF1- and ORF2-encoding polyproteins, which are nearly identical, and concluded as the structural proteins.

### DtBRV1 is efficiently transmitted vertically while not horizontally

The conidia of strain CJP4-1 were promptly produced by inoculation of the mycelia on alfalfa sticks, which were incubated in PDA plates, and the resulting conidia were collected at 15 day post-inoculation (dpi), diluted and cultured on PDA plates for colony formation. A total of 23 conidium-generating subisolates (termed CJP41-D1 to D23) were picked and transferred into new PDA plates for viral detection by dsRNA extraction. The results showed that 18 out of 23 isolates (accounting for 78.3%) had DtBRV1 dsRNAs in their nucleic acid preparations, while the remaning five isolates (CJP4-1D7, -D8, -D10, -D12, and -D14) did not (Fig. 4A). This was ratified following RT-PCR detection of dsRNA 2 using a pair of oligonucleotide primers DtBRV1-F1/-R1 (Fig. 4B and Table S1).

To investigate the horizontal transmission of DtBRV1, DtBRV1-infected strain CJP4-1 (donor) and DtBRV1-free strain JYC1-6 (recipient) of *D. theifolia* were dual-cultured for ten days (Fig. S3A). A total of fifteen (with three from each of five dual-cultures) JYC1-6 mycelium discs far from the contact area were excised, transferred into new PDA plates for colony formation, and analyzed for the presence of DtBRV1 dsRNAs. None (0/15) of the subisolates were infected by DtBRV1, it might be due to an antagonistic effect between the mycelia of both strains (Fig. S3A). Therefore, strain CJP4-1 was further dual-cultured with another DtBRV1-free strain *D. theifolia* strain JYC1-9 (free of DtBRV1), without an antagonistic band being observed (Fig. S3B), and fifteen JYC1-9 subisolates were analyzed, resulting in no subisolates positive for DtBRV1 as well. These results indicated that DtBRV1 is difficult to be horizontally transmitted to other *D. theifolia* strains through hyphal fusion.

### Protoplasts of strain JYC1-6 are transfected successfully with purified virions

To obtain another *D. theifolia* strain carrying DtBRV1, we attempted to transfect the protoplasts of strain JYC1-6 with purified DtBRV1 particles in the presence of PEG 6000 due to the failure of the dual-culture experiment. A total of 107 derivative isolates were picked from the generated protoplasts of strain JYC1-6 and transferred into new PDA plates for viral detection by dsRNA extraction. The results showed that four isolates (JYC1-6T43, -T49, -T50, and -T88), accounting for 2.8% incidence, had DtBRV1 dsRNAs in their nucleic acid preparations (Fig. 5A). This was ratified following RT-PCR detection of dsRNA 2 using a pair of oligonucleotide primers DtBRV1-F1/-R1 (Fig. 5B and Table S1).

**Fig. 5.**
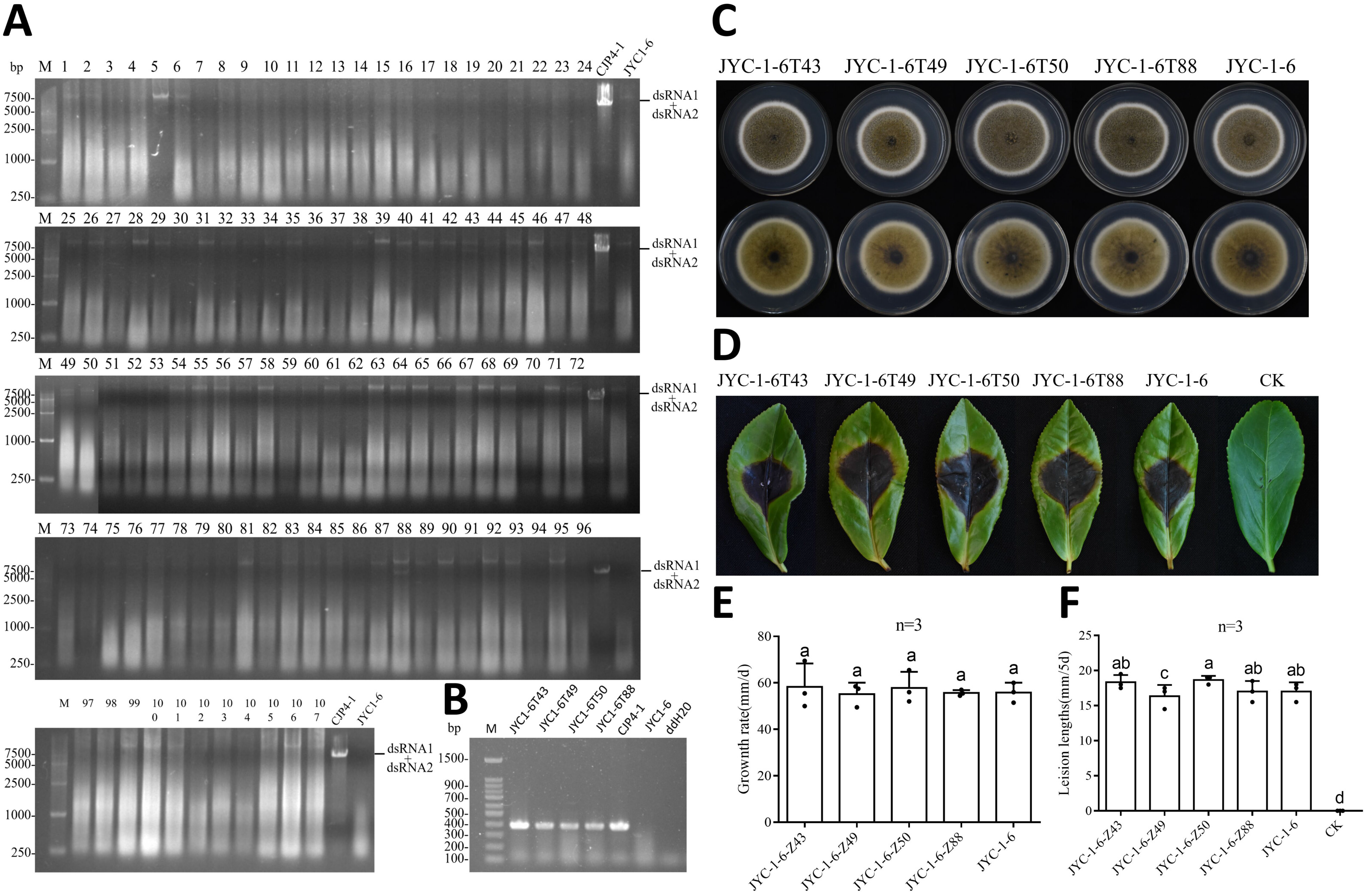
Effects of DtBRV1 on fungal morphology, growth, and pathogenicity. (A) DsRNA extraction from the mycelia of generated protoplasts of strain JYC-1-6 was transfected with DtBRV1 virions. Lanes 1 to 107 are transfectants (JYC1-6T1 to JYC1-6T107). CJP-4-1 was used as a positive control and JYC1-6 as a negative control. (B) RT-PCR identification of the DtBRV1 absence in transfectants (JYC1-6-Z43, -T49, -T50, and -T88), with strain CJP4-1 involved as a positive control resulting in the target product in size of 390 bp. JYC1-6 was used as a negative control and ddH_2_O as blank control. (C) Colony morphologies of DtBRV1-infected transfectants JYC1-6T43, -T49, -T50, -T88 and DtBRV1-free strain JYC1-6 cultured on PDA at 25°C for 8 days. (D) Virulence assessment of strain JYC1-6 and its transfectants on tea leaves (*C. sinensis* var. E’cha no.1). CK, the black control inoculated with uncolonized PDA plugs. (E and F) Bar graphs of the growth rates (E) and lesion lengths (F) of strain JYC1-6 and its transfectants.

### DtBRV1 is association with the fungal morpoglogical changes while not with the attenuated virulence

To check whether DtBRV1 has some effects on the fungal biological traits, the parent strain CJP4-1 together with its subisolates free of DtBRV1 including CJP4-1D7, -D8, -D10, -D12, and -D14 were cultured on PDA plates to access their morphologies and growth rates (Fig. 4C and 4E). The results showed that all these virus-free isolates grew at similar growth rates but with more dense aerial mycelia as compared with those of the parent strain CJP4-1. As inoculated on detached tea leaves (*C. sinensis* var. E’cha no.1), all these virus-free subisolates, similar to strain CJP4-1, induced no obvious lesions (Fig. 4D and 4F). To further clarify whether DtBRV1 is responsible for the morphological changes and the avirulence, the DtBRV1-transfected subisolates (JYC1-6-1T43, -T49, -T50, and -T88) together with the parent strain were further accessed their morphologies, growth rates, and virulence, and all these subisolates had growth rates and lesion lengths on the tea leaves similar to those of the parent strain (Fig. 5C to 5F), suggesting that DtBRV1 has no or undetectable effects to the biological traits of the hots fungi.

## DISCUSSION

In this study, three dsRNAs (dsRNAs 1 to 3) were detected from an avirulent strain CJP4-1 of *D. theifolia* and two (1 and 2) were identified as the genomic components of a novel botybirnavirus, named DtBRV1, based on sequence alignment, ultracentrifugation, TEM observation, and phylogenetic analysis. DtBRV1 possesses some molecular traits typical for other members under this genus: 1) it has isometric viral particles in ∼39.8 nm, fitting in the diameter range of 35-40 nm for other botybirnaviruses; 2) its genome comprises two segments in similar sizes; 3) each genomic segment contains a single large ORF, with an RdRp domain and a proline-rich region in one of them; 4) its dsRNAs have a long and conserved 5’-UTR and a short 3’-UTR. Moreover, DtBRV1 dsRNAs 1 and dsRNA2 share a high identity of 42.12%, with a long (433 bp) and conserved 5’-UTR, and both ORFs initiate at the end of the conserved terminal sequence (Fig. 2B). This molecular trait with long and strictly conserved terminal sequences has been observed in other botybirnaviruses and Rosellinia necatrix megabirnavirus 1 (RnMBVl), but it is rarely observed in other bipartite and multicomponent dsRNA viruses (22). It is worth noting that a small dsRNA fragment (dsRNA3) in size of 727 bp was detected in association with DtBRV1, and shares a considerably high identity (approximately 40%) with the left regions of dsRNAs 1 or 2, and it suggests that dsRNAs is not the satellite of DtBRV1 since a satellite always shares no similarity with the genome of its host virus, together with that dsRNAs could not co-precipitated with dsRNAs 1 and 2 by ultracentrifugation (Fig. 3A). A 1.7 kb-sized dsRNA fragment was ever discovered co-infecting with SsBRV1 in *S. sclerotiorum*, and was concluded as a satellite-like RNA since it could be co-precipitated together with SsBRV1(18). Therefore, dsRNA3 is unlike the satellite-like RNA of SsBRV1, and how it could survive and replicate in vivo require further studies.

Five fungal botybirnaviruses have been reported, and all infected filamentous Ascomycota fungi, including SsBRV1 and SsBRV2 from *S. sclerotiorum* (18, 19), BpRV1 from *B. porri*(17), and ABV1 and AaBbV1 from *Alternaria* fungi(20, 21). Besides, an unreported member designated as Bipolaris maydis botybirnavirus 1 (BmBRV1, accession no. MF034087.1 and MF034086.1) from *B. maydis* has been deposited in NCBI, and Soybean leaf-associated botybirnavirus 1 (SlaBRVl), which was discovered from metatranscriptomics survey of soybean phyllosphere phytobiomes(23). To our knowledge, this is the first report of a botybirnavirus infecting *D. theifolia*.

DtBRVl is encapsidated in isometric viral particles, two major protein bands (p65 and p75) were observed by SDS-PAGE, and we concluded that p65 and p75 are two major structural proteins composing DtBRVl coat proteins (CPs). By PMF-MS analysis, p65- and p75-generating peptides are nearly identical, and matched partial sequences of both ORF1- and ORF2-encoding proteins, dominantly in their left regions and mostly apart from the RdRp domain, suggesting that p75 (or p65) is a fused protein derived from both left regions of ORF1- and ORF2-encoding polyproteins after some processing steps including cleavage and linkage. CP generating from a large polyprotein through cleavage is commonly observed in viruses that have a large genome encoding a polyprotein, e.g., Ustilago maydis virus (24) and bursal disease virus (25, 26). Two structural proteins are nearly identical, which has been observed in other botybirnaviruses, e.g. for BpRV1, SsBRV1, and SsBRV2, and this phenomenon has been considered that the small protein possibly degraded from the large one, or derived from a large molecule protein *via* a posttranslational process, similar to Helmintllosporium victoriae virus 190S (17-19, 27). A major protein (p65 or p75) suggests an icosahedral T=1 capsid consisting of 60 copies of one single polypeptide. This situation is similar to some typical chrysoviruses, belonging to the neighboring family in the phylogenetic tree (Fig. 2D), e.g., Penicillium chrysogenum virus (PcV) (28) and Cryphonectria nitschkei virus 1 (CnV1) (29), but distinct from those of chrysoviruses including Botryosphaeria dothidea chrysovirus 1 (BdCV1), Botrytis porri RNA virus 1 (BpRV1) (17) and quadriviruses (30), reported to have a T=1 capsid made up by 60 CP heterodimers, and Magnaporthe oryzae chrysovirus 1 (MOCV1) with multiple CP components (31, 32).

To clarify the biological traits related to DtBRV1 infection, vertical and horizontal transmission experiments were accessed with conidium-generating subisolates of strain CJP4-1 and dual-culture with other *D. theifolia* strains, suggesting that DtBRV1 is efficiently vertically transmitted while difficult in horizontal transmission from strain CJP4-1 to other strains, which might be due to vegetative incompatibility between these strains. As DtBRV1 was eliminated from strain CJP4-1, the derived subisolates grew at similar growth rates but with more dense mycelia as compared with the parent strain, and still kept avirulent as accessed their virulence on tea leaves. As transfection of purified DtBRV1 virions into virulent strain JYC1-6, it did not cause a reduction in mycelial growth and hypovirulence of the transfectants. These results suggest that DtBRV1 most likely has no or tiny effects on the growth and virulence of the host fungus. Therefore, we speculate that the avirulent traits of CJP4-1 are not related to DtBRV1 infection. Similarly, AaBbV1 and SsBRV1 had no significant effect on the host fungi (18, 20), although BpRV1 and SsBRV2 infection attenuate the virulence of the host fungi (17, 19).

Most mycoviruses have been largely isolated and characterized from phytopathogenetic fungi infecting cereal and oil crops, fruit and forest trees, while rarely from tea plants, an important economic crop. Here we present the first report of a botybirnavirus, as well as the first mycovirus specimen, infecting *D. theifolia*, which would be expected to contribute to a better understanding of the viral community in association with the phytopathogenetic fungi infecting tea plants.

## MATERIALS AND METHODS

### Fungus strains

Strains CJP4-1, JYC1-6, and JYC1-9 of *D. theifolia* strains were isolated from tea leaves with brown spot symptoms that were collected from Zigui county, Hubei province, China, and identified by combining sequence data of internal transcribed spacer (ITS), partial β-tubulin (TUB), partial RNA polymerase II second largest subunit (RPB2), partial large subunit ribosomal RNA (LUS) gene regions. Of them, strains JYC1-6 and JYC1-9 (previously termed JYC-1-6 and JYC-1-9, respectively) have been characterized as the etiology of tea leaf brown-black spot disease (16).

### DsRNA extraction

Fungal dsRNA was extracted using a silica spin column-based method as previously described (33). Fungal mycelium discs (5 mm in diameter) were cultured on PDA plates covered with sterilized cellophane for 5 days. Approximately 0.3 g of fresh mycelium was ground into powder in liquid nitrogen for dsRNA extraction. dsRNA was treated by enzymatic digestion with DNaseI and S1 nuclease (New England Biolabs) to remove DNA and ssRNA from the samples. The resulting dsRNA was fractionated by agarose gel (1%, w/v) electrophoresis, stained with ethidium bromide (0.1 mg/mL), and detected by UV transillumination.

### NGS sequencing, terminal sequence determination, and RT-PCR amplication

The mycelia of strain CJP4-1 were collected and subjected to long non-coding RNA sequencing (lncRNA-seq) in Personal Biotechnology Co., Ltd. (Shanghai, China) by Illumina sequencing platform (Illumina Novaseq 6000). The resulting raw data were trimmed and cleaned by removing low-quality tags, polyA or N tags, and assembled using IDBA-UD (34). Clean reads were assembled *de novo* using Velvet with a k-mer value of 17 (35) and CAP3 with default values (36), and the contigs were assembled in Vector NTI Advance (version 11.5) software.

Terminal cDNA sequences of dsRNAs were obtained by RACE protocols as previously described (37). Briefly, the 3′ terminus of each strand of dsRNA was ligated with PC3-T7loop oligo (Table S1) using T4 RNA ligase (TaKaRa, China) at 16 °C for 16 h. The oligonucleotide-ligated dsRNA was purified, denatured in DMSO, subsequently reverse transcribed with M-MLV reverse transcriptase and 3 pmol of PC2, a primer complementary to the oligonucleotide used for the RNA ligation (Table S1), and subjected to PCR amplification in combination with the specific primers (Table S1) homology (for 3′-RACE) or complementary (5′-RACE) to the obtained sequences, respectively. The PCR amplicons were purified, and cloned into pMD18-T vector (TaKaRa, China) and the recombinant plasmids were picked and sent to Tsingke Biotechnology Co., Ltd. (Wuhan, China) for sequencing. Every nucleotide was determined with at least three independent overlapping clones in both orientations.

### Phylogenetic and sequence analyses

The full-length cDNA sequences of the dsRNAs were used to predict the possible ORFs and the corresponding amino acid sequences with ORF finder program (https://www.ncbi.nlm.nih.gov/orffinder/). The conserved structural domains of viral proteins were predicted on the PFAM website (http://genome.jp/tools/motif/). The sequence search and homologous matching of the fungal viruses were performed in the NCBI database, and the protein sequences of the fungal viruses with high similarity and some other dsRNA viruses were imported into PhyloSuite (version 1.2.2) software for subsequent multiple sequence matching and phylogenetic analysis (38). Multiple alignments of nucleic acid or amino acid sequences were performed with MAFFT (version 7.313) software, and the obtained data were distinguished from each other with bright colors in GeneDoc software (http://www.nrbsc.org/downloads). Maximum likelihood phylogenies were inferred using IQ-TREE (39) under the model automatically selected by IQ-TREE (‘Auto’ option in IQ-TREE) for 1000 ultrafast (40) bootstraps, as well as the Shimodaira–Hasegawa–like approximate likelihood-ratio test (41). Visualization and refinement of the phylogenetic tree on the ITOL online site(https://itol.embl.de/) (42).

### Virus particles purification and polypeptide mass fingerprint-mass spectrum (PMF-MS) analysis

The isolation and purification of viral particles were performed as previously described (13) with some modifications. Briefly, a total of 36 g of mycelia were harvested and ground with liquid nitrogen after grown on sterilized cellophane placed on PDA at 25°C for 7 days, transferred into new tubes containing 3 volumes of 0.1 mol/L phosphate buffer (PB, 8.0 mmol/L Na_2_HPO_4_, 2.0 mmol/L NaH_2_PO_4_, pH 7.0) and 0.5%(V/V) β-mercaptoethanol, mixed gently and centrifuged at 12,000 rpm and 4°C for 15 min, and subsequently subjected to ultracentrifugation at 32000 rpm for 3 h for two times (Optima LE-80K, Backman Coulter, Inc. USA). The resultant sediment was resuspended in 0.1 mol/L PB and centrifuged in sucrose density gradients of 10–50% (W/V) with an increment of 10% at 32000 rpm for 3 h. Following sucrose gradient ultracentrifugation, each sucrose fraction was collected, diluted, and subjected to ultracentrifugation to pellet virus particles, and resuspended in 150 μL of 0.1 mol/L PB. Each gradient of virus suspensions was extracted using phenol and chloroform and analyzed by agarose gel electrophoresis (1%, W/V) to detect viral dsRNA. Moreover, each suspension was adsorbed on the copper grids for 3 min, placed on filter paper to dry for 2 min, stained with 2% (W/V) phosphotungstic acid (PTA) for 3 min in a drop of 2% (W/V) phosphotungstic acid (PTA), and observed with a transmission electron microscope(TEM, H7650; Hitachi). Virion sizes were measured with ImageJ software.

A total of 200 μL suspension from each sucrose fraction was subjected to protein extraction, analyzed by 12% SDS-PAGE with 25 mM Tris/glycine and 0.1% SDS, and stained with Coomassie brilliant blue R-250 (Bio-Safe CBB; Bio-Rad, USA). Protein bands on the gel were individually excised and subjected to PMF analysis at Sangon Biotech (Shanghai) Co., Ltd, China.

### Vertical and horizontal transmission of mycovirus

Vertical transmission of a mycovirus was accessed *via* conidia as previously described (43). Fungal mycelia were inoculated on alfalfa sticks and incubated in PDA plates for 15 days for conidium formation at 25°C in darkness. The resulting conidia were diluted into the concentrations of 10^4^-10^5^ conidia/mL, and 100 μL of the conidial suspension were smeared onto PDA media and cultured at 25°C in darkness for colony formation. The conidium-generating colonies were transferred into new PDA plates for further growth and dsRNA extraction.

Horizontal transmission of a mycovirus was accessed *via* hyphal anastomosis as previously described (13). Briefly, a donor strain was dually cultured with a recipient strain (or subisolate) at 25°C for 7-10 days to allow the two colonies to contact each other in each PDA plate. After incubation of the contact cultures, mycelial agar plugs were excised from the colony margin of the recipient strain (apart from the donor colony as far as possible) and transferred onto a new PDA plate for colony formation for dsRNA extraction. Each contact culture was conducted in triplication, five derived isolates from each recipient strain and fifteen mycelium plugs in total were obtained in the contact cultures.

### Protoplast transfection with DtBRV1 virions

Protoplasts were isolated and transfected with purified viral particles as previously described (44). Protoplasts were isolated from actively growing mycelia of the DtBRV1-free *D. theifolia* strain JYC1-6, and transfection with 70.0–80.0 μg DtBRV1 virions in the presence of PEG 6000. The transfected protoplast suspension was incubated without shaking at 25°C in the dark for approximately 16 hours and spread onto BA (bottom agar) plates containing 0.3% yeast extract, 0.3% casein hydrolysate, 20% sucrose, and 1% agar). Mycelial agar plugs generated from the protoplasts were transferred into new PDA plates for further growth and dsRNA extraction.

### Growth rate, morphology, and virulence assays

Fungal growth rates and morphologies were assessed in triplication for each strain or subisolate as previously described (13). Fungal virulence was determined in three replication for each strain or subisolate following inoculation of detached tea leaves (*C. sinensis* var. E’cha no.1) under wounded conditions as previously described (13). At 5 dpi, lesions that developed on the inoculated leaves were taken photos and measured.

## Data availability statement

The complete nucleotide sequence of DtBRV1 reported in this paper is deposited in the GenBank™ database with the accession number XXX to XXX.

## Author contributions

W.X. supervised and designed the experiments and revised the manuscript; L.Y. conducted most of the experiments and wrote the manuscript; X.S. conducted part of the biological tests, Y.H. isolated fungal strains, and J.C. and Q.X were involved in part of the experiments; K.S. improved the English. All authors have read and agreed to the published version of the manuscript.

## Acknowledgments

All the authors thank the financial support by the National Science Foundation of China and Hubei Hongshan Laboratory.

